# Genome-Wide CRISPR Screening Identifies BRD9 as a Druggable Component of Interferon-Stimulated Gene Expression and Antiviral Activity

**DOI:** 10.1101/2021.02.04.429732

**Authors:** Jacob Börold, Davide Eletto, Idoia Busnadiego, Nina K. Mair, Eva Moritz, Samira Schiefer, Nora Schmidt, Philipp P. Petric, W. Wei-Lynn Wong, Martin Schwemmle, Benjamin G. Hale

**Affiliations:** Institute of Medical Virology, University of Zurich, Zurich, Switzerland; Life Science Zurich Graduate School, ETH and University of Zurich, Zurich, Switzerland; Institute of Virology, Freiburg University Medical Center, Faculty of Medicine, University of Freiburg, Freiburg, Germany; Spemann Graduate School of Biology and Medicine, University of Freiburg, Freiburg, Germany; Institute of Experimental Immunology, University of Zurich, Zurich, Switzerland

## Abstract

Transient interferon (IFN) induction of IFN-stimulated genes (ISGs) creates a formidable protective antiviral state. However, loss of appropriate control mechanisms can result in constitutive pathogenic ISG upregulation. Here, we used genome-wide loss-of-function screening to establish genes critical for IFN signaling, identifying all expected members of the JAK-STAT pathway and the previously unappreciated bromodomain-containing protein 9 (BRD9), a defining subunit of non-canonical BAF (ncBAF) chromatin remodeling complexes. Genetic knock-out or small-molecule mediated degradation of BRD9 limited IFN-induced expression of a subset of ISGs in multiple cell-types, and prevented IFN from exerting full antiviral activity against several RNA and DNA viruses. Mechanistically, BRD9 acts at the level of ISG transcription, exhibits a proximal association with STAT2 following IFN stimulation, and relies on its intact acetyl-binding bromodomain and unique ncBAF scaffolding function for activity. Given its druggability, BRD9 may be an attractive target for dampening constitutive ISG expression under certain pathogenic autoinflammatory conditions.

## INTRODUCTION

The type I interferon (IFN-α/β) system is a key component of the human innate immune response, and acts as a first line of defense against invading pathogens such as viruses. At a general level, following host recognition of infection, secreted type I IFNs relay signals to target cells via their binding to cognate cell surface receptors (IFNAR1/IFNAR2) and the subsequent triggering of a well-defined phosphorylation-activation JAK-STAT signaling cascade involving the Janus kinases (JAK1/TYK2) and Signal Transducer and Activator of Transcription (STAT1/STAT2) proteins. Together with IFN-regulatory factor 9 (IRF9), activated STAT1 and STAT2 form a heterotrimeric transcription factor complex, termed ISGF3, that facilitates the transcription of hundreds of antiviral IFN-stimulated genes (ISGs)^1–3^. The critical nature of the IFN system in protecting humans against infection is exemplified by findings that some individuals with genetic defects in IFN pathway components can exhibit severe, or even fatal, viral diseases, particularly in the absence of pre-existing humoral immunity, as may be the case in young infants or following infection with an antigenically-novel pandemic virus^4,5^.

It is essential that the IFN system is tightly regulated at the molecular level to prevent exuberant proinflammatory responses following infection^6–8^. Furthermore, uncontrolled activation of the IFN system caused by loss- or gain- of-function mutations in key regulators of the IFN pathway can be associated with aberrantly high levels of circulating IFNs and/or the constitutive expression of ISGs, leading to a broad range of autoinflammatory disorders known as interferonopathies^9–17^. Proposed treatments for interferonopathies include JAK inhibitors (such as baricitinib^18^, ruxolitinib^19^ or tofacitinib^20^) to limit the signaling action of high levels of constitutively circulating IFN or inappropriate JAK-STAT regulation. However, while effective in treating interferonopathies, inhibiting critical components of the IFN signaling pathway can increase patient susceptibility to some viral infections^21,22^. Therefore a nuanced approach targeting factors involved in transcription of specific ISGs may be more appropriate^22^.

In this study, we sought to leverage CRISPR/Cas9-mediated genome-wide loss-of-function screening to interrogate the IFN signaling cascade and identify human host factors required for ISG expression. Our screening results led us to focus on Bromodomain-containing protein 9 (BRD9), a defining subunit of specific chromatin-remodeling complexes^23–25^, which has not previously been implicated in IFN responses. Many factors involved in transcriptional regulation via chromatin modification or remodeling play important roles in STAT-mediated ISG expression^22,26–39^, and some (such as histone deacetylases (HDACs) and bromodomain-containing proteins^22,26,35,39^) can be targeted pharmacologically. Here, we provide evidence that BRD9 contributes to the expression of a subset of ISGs (and thus the full antiviral action of IFN) via its unique functions in non-canonical BAF (BRG1- or BRM- associated factor; ncBAF) chromatin remodeling complexes. Importantly, this contribution of BRD9 is conserved across numerous human cell types, including primary cells, and against multiple distinct viruses. Moreover, BRD9 can be specifically targeted for degradation by small-molecule compounds without impacting cell viability. Our data add mechanistic insights into human factors required for the full activity of IFN, and suggest a plausible therapeutic target to rationally temper detrimentally-high ISG levels in some autoinflammatory disorders, including interferonopathies.

## RESULTS

### A Genome-Wide Loss-of-Function Screen Identifies Human Genes Important for Interferon-Stimulated Gene Expression

The type I IFN signaling pathway leading to induction of ISG expression is comprised of several canonical gene components, including receptors (*IFNAR1*, *IFNAR2*), kinases (*JAK1*, *TYK2*), and transcription factors (ISGF3: *STAT1*, *STAT2*, *IRF9*) (**Figure 1A**). To identify additional, previously uncharacterized, genes critical for the ability of IFN to signal and induce ISG expression, we performed a genome-wide FACS-based loss-of-function screen in the human lung epithelial cell-line, A549. We used the GeCKOv2 CRISPR-Cas9 library, which contains a total of 123,411 CRISPR single-guide RNAs (sgRNAs) targeting 19,050 genes in the human genome with 6 guide RNAs per gene^40^. First, we generated a sub-clone of an A549-based reporter cell-line that expresses eGFP under control of an IFN-stimulated response element (A549/pr(ISRE).eGFP.A1)^41^, and confirmed that treatment of this sub-clone with IFN-α2 leads to high and homogenous expression of eGFP, with clear separation of IFN-stimulated and non-stimulated cell populations in flow cytometry analysis (**Figure 1B**). As a pre-screen validation, we next transduced this reporter cell-line with all-in-one puromycin-resistant lentiviral vectors expressing Cas9 and several control sgRNAs selected from the GeCKOv2 human library. Following puromycin selection for 10 days to ensure protein depletion after CRISPR gene editing, we observed that a significant proportion of reporter cells transduced with sgRNAs targeting the canonical pathway components, *IFNAR1* or *STAT1*, failed to express eGFP in response to IFN-α2 stimulation (**Figure 1C**). As expected, reporter cells transduced with an sgRNA targeting a non-pathway component, *IFNLR1*, responded similarly to IFN-α2 stimulation as the parental cell-line (**Figure 1C**). These data indicate that a FACS-based loss-of-function screen based on the GeCKOv2 human library can be used to identify components of the type I IFN signaling pathway in A549 cells.

**Figure 1.**
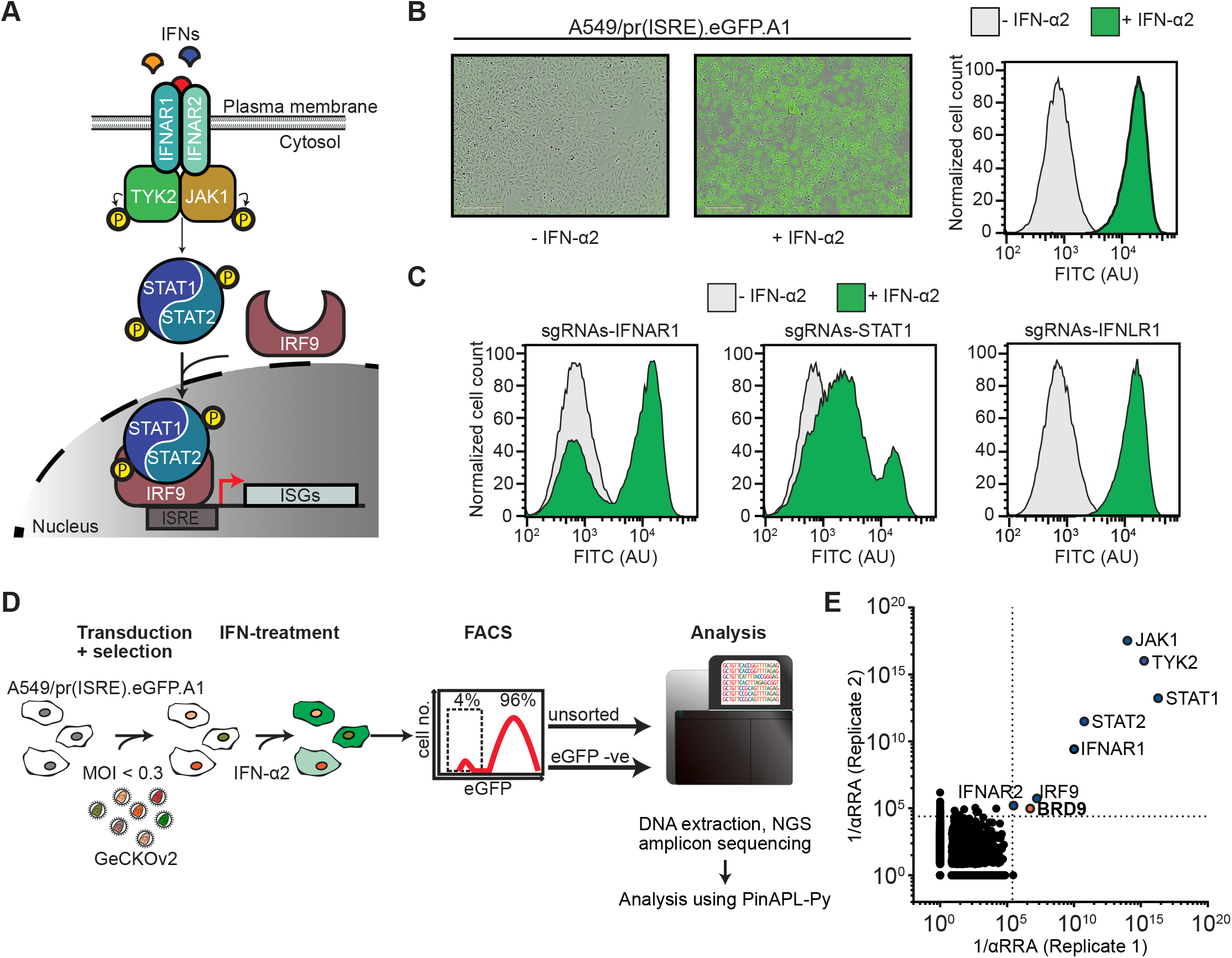
A Genome-Scale Loss-of-Function Screen Identifies Human Genes Important for Interferon-Stimulated Gene Expression. **(A)** Schematic representation of the type I IFN signaling pathway in human cells. **(B)** Stimulation of A549/pr(ISRE).eGFP.A1 cells with 1000 IU/mL of IFN-α2 for 16h results in high and homogeneous expression of eGFP as determined by fluorescence microscopy (left panels), and flow cytometry (right panel). The FITC channel was used to monitor eGFP levels by flow cytometry, and AU refers to arbitrary units. Data are representative of at least two independent replicates. **(C)** A549/pr(ISRE).eGFP.A1 cells were transduced for at least 10 days with lentivirus pools expressing Cas9 and sgRNAs targeting the indicated host genes. eGFP levels following 16h of IFN-α2 treatment (1000 IU/mL) or mock, were determined by flow cytometry. Data are representative of at least two independent replicates. **(D)** Workflow overview of the genome-scale loss-of-function screen for essential IFN signaling components using the full GeCKOv2 CRISPR-Cas9 library. **(E)** 1/αRRA scores from two independent biological replicates of the genome-scale screen. Dotted lines represent the significance cut-offs for each replicate. Genes identified as significant in both screens are highlighted. See also **Supplementary Figure 1** and **Supplementary Dataset 1**.

We transduced A549/pr(ISRE).eGFP.A1 cells with a pool of all-in-one puromycin-resistant lentiviral vectors (lentiCRISPRv2) comprising the entire GeCKOv2 human library at a multiplicity of infection (MOI) of 0.3 focus forming units (FFU)/cell to ensure that most transduced cells received only one sgRNA. We then selected the cells with puromycin for 10 days to ensure that only transduced cells survived, and to allow for protein depletion after genome editing. Puromycin-selected reporter cells were then treated (or not) with 1000 IU/mL IFN-α2 for 16h to stimulate eGFP expression, and FACS was used to enrich for eGFP negative cells that failed to respond to IFN-α2 (**Figure 1D**). Following extraction of genomic DNA from both the parental library-transduced reporter cell-line and the eGFP negative cells that failed to respond to IFN treatment, sgRNA abundance was determined via targeted sequencing (**Figure 1D**). Two independent biological replicates of the screen were performed, and for each replicate the adjusted robust-rank aggregation (αRRA) feature of the PinAPL-Py platform was used to determine gene-level rankings (based on relative enrichment of each individual sgRNA^42,43^) for loss-of-function events leading to IFN non-responsiveness. From this genome-wide screen, we identified 8 genes that exhibited significant αRRA enrichment (p < 0.01) in both independent replicates (**Figure 1E & Supplementary Dataset 1**). The robustness of our results was validated by the observation that 7 of these 8 genes comprise all the known essential factors in the canonical type I IFN signaling pathway leading to ISG expression: *IFNAR1*, *IFNAR2*, *JAK1*, *TYK2*, *STAT1*, *STAT2*, and *IRF9*. The 8^th^ gene found to be significantly enriched in both our screens was *BRD9*, which has not previously been characterized for its role in the type I IFN signaling pathway. To validate this result, we performed independent experiments using individually-cloned and arrayed sgRNAs, and could confirm that all 6 *BRD9*-targeting sgRNAs within the GeCKOv2 human library limit IFN-α2-stimulated expression of eGFP in A549/pr(ISRE).eGFP.A1 cells (**Supplementary Figure 1A**). Notably, a recent independent genome-wide screening approach using insertional mutagenesis in HAP1 cells also identified *BRD9* enrichment when searching for positive regulators of type III IFN-stimulated gene expression^8^, which is thought to use the same intracellular signaling machinery as type I IFN^1^. Thus, our genome-wide loss-of-function screen successfully identified both known genetic components of the human type I IFN signaling pathway, and a new uncharacterized factor, *BRD9*.

### BRD9 is Important for Interferon-Stimulated Gene Transcription and Antiviral Activity

*BRD9* encodes Bromodomain-containing protein 9 (BRD9). BRD9 is a defining subunit of the recently described non-canonical BAF (ncBAF) complex, one of three major distinct types of ATP-dependent chromatin remodeling complexes (together with canonical BAF and the polybromo-associated, PBAF) that govern DNA accessibility and thus appropriate regulation of gene transcriptional programs (**Supplementary Figure 1B**)^23–25^. To dissect the functions of BRD9 in the human type I IFN signaling pathway, we used CRISPR/Cas9 to generate two independent sets of A549-based BRD9 knock-out (KO) and control (CTRL) clonal cell-lines. Amplicon sequencing of genomic DNA confirmed the KO and CTRL genotypes of all clones, and western blot validated the loss of BRD9 protein expression in both KO clones (**Figures 2A-B & Supplementary Figures 2A-B**). Loss of BRD9 did not interfere with IFN-α2-induced phosphorylation of JAK1 or STAT1, or with the IFN-α2-induced translocation of phosphorylated STAT1 to the nucleus (**Figures 2C-D**). However, consistent with its known role in regulation of chromatin remodeling and gene transcription, loss of BRD9 led to a reduction in IFN-α2-induced *MX1* mRNA levels and a consequent reduction in IFN-α2-induced MxA protein levels (**Figures 2E-F & Supplementary Figure 2C**). Importantly, loss of BRD9 rendered cells unable to mount an effective IFN-α2-stimulated antiviral response against influenza A virus (IAV) (**Figure 2G & Supplementary Figure 2D**). To confirm that the observed phenotypes were due only to the loss of BRD9 (and not to off-target CRISPR edits), we used a lentivirus vector to reconstitute BRD9 expression. Reconstitution of BRD9 in BRD9-KO cells fully restored the ability of IFN-α2 to induce MxA protein levels (**Figure 2H & Supplementary Figure 2E**), as well as the ability of IFN-α2 to mediate antiviral activity against IAV (**Figure 2I & Supplementary Figure 2F**). These results identify a role for BRD9 in type I IFN signaling at the level of ISG transcription, and reveal that this role is critical for the effective antiviral activity of IFN.

**Figure 2.**
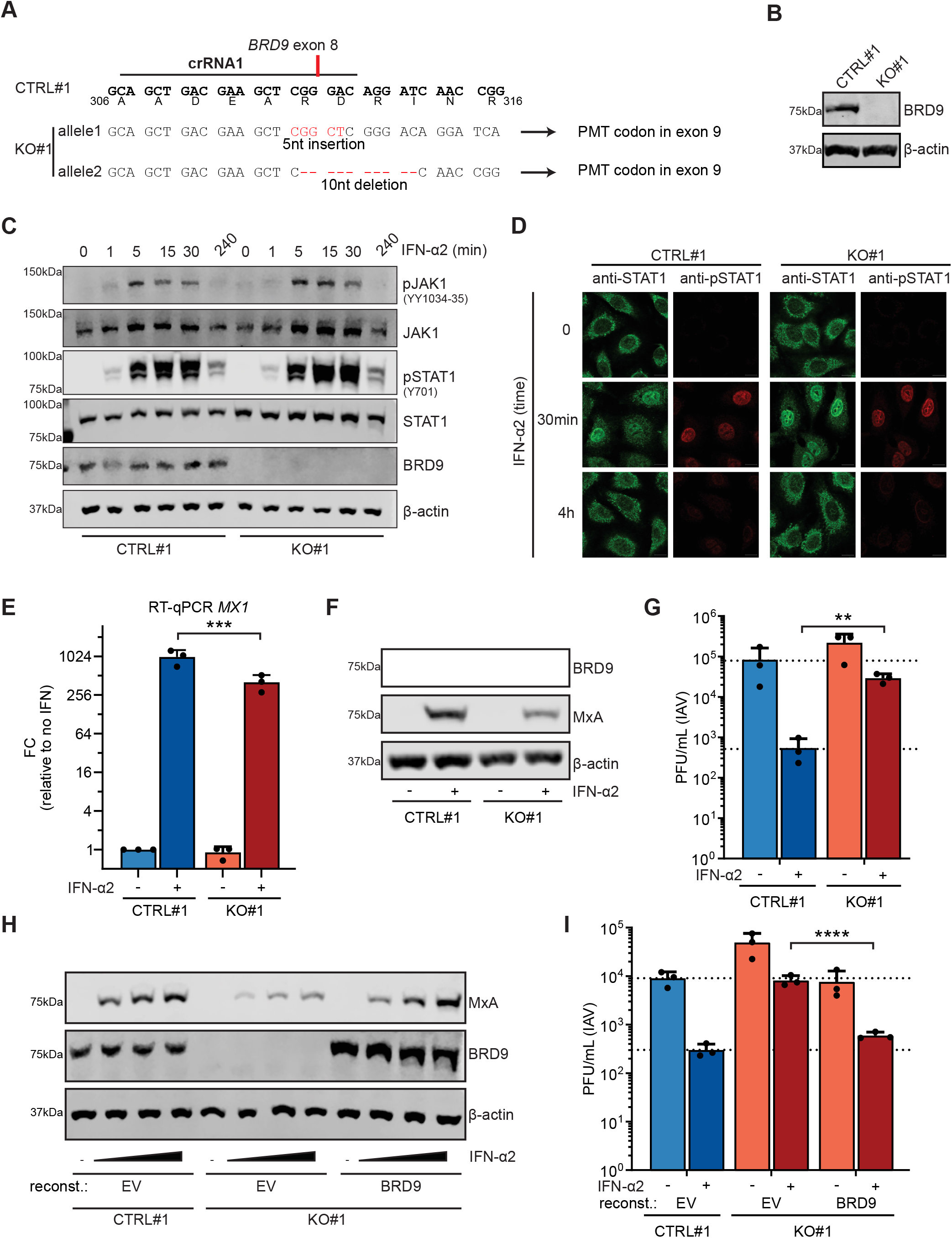
BRD9 is Important for Interferon-Stimulated Gene Transcription and Antiviral Activity. **(A)** An A549-derived BRD9-KO cell-clone (KO#1) was generated using a crRNA targeting exon 8 of *BRD9*. The target sequence of the crRNA (termed crRNA1), and the resulting genomic alterations determined by NGS for the two *BRD9* alleles, are shown in comparison to an unedited control clone (CTRL#1). Generated indels lead to a premature termination codon (PMT) in the following exon. Encoded amino-acids are shown below the CTRL nucleotide sequence (bold indicates nucleotides in exons). **(B)** Western blot analysis of lysates from CTRL#1 or BRD9-KO#1 cells. BRD9 and β-actin were detected with specific antibodies. Data are representative of at least two independent replicates. **(C)** Western blot analysis of CTRL#1 or BRD9-KO#1 lysates from cells treated for the indicated times with 1000 IU/mL of IFN-α2. The indicated proteins were detected with specific antibodies. Data are representative of at least two independent replicates. **(D)** Immunofluorescence analysis of CTRL#1 or BRD9-KO#1 cells treated for the indicated times with 1000 IU/mL of IFN-α2. The indicated proteins were detected with specific antibodies. Data are representative of at least two independent replicates. **(E)** RT-qPCR analysis of *MX1* levels in CTRL#1 or BRD9-KO#1 cells following treatment, or not, with 100 IU/mL of IFN-α2 for 6h. *GAPDH* transcript levels were used for normalization. Data represent means and standard deviations from three independent experiments (individual data points shown). Statistical significance was determined by unpaired 2-tailed t-test on ΔC_t_ values (***p-value < 0.001). **(F)** Western blot analysis of CTRL#1 or BRD9-KO#1 lysates from cells treated or not for 16h with 1000 IU/mL of IFN-α2. The indicated proteins were detected with specific antibodies. Data are representative of at least two independent replicates. **(G)** CTRL#1 or BRD9-KO#1 cells were treated, or not, with 1000 IU/mL of IFN-α2 for 16h prior to infection with IAV (WSN/33) at an MOI of 0.01 PFU/cell. Viral titers were determined after 24h by plaque assay. **(H)** CTRL#1 or BRD9-KO#1 cells were stably-transduced with BRD9-expressing, or control (EV, empty vector), lentiviruses and treated with a range of IFN-α2 concentrations (0, 10, 100, 1000 IU/mL) for 16h prior to lysis and analysis for the indicated proteins by western blot. Data are representative of at least two independent replicates. **(I)** CTRL#1 or BRD9-KO#1 cells were stably-transduced with BRD9-expressing, or control (EV, empty vector), lentiviruses and treated with 1000 IU/mL of IFN-α2 for 16h prior to infection with IAV (WSN/33) at an MOI of 0.01 PFU/cell. Viral titers were determined after 24h by plaque assay. For (G) and (I), data represent means and standard deviations from three independent experiments (individual data points shown). Statistical significance was determined by 1-way ANOVA on log-transformed plaque counts (**p-value < 0.001; ****p-value < 0.0001). See also **Supplementary Figure 2**.

### Targeted Degradation of BRD9 Reveals its Cell-Type Independent Contribution to Interferon-Stimulated Antiviral Activity Against Multiple Viruses

We used a targeted chemical degrader of BRD9, dBRD9-A, as a tool to efficiently deplete cells of BRD9 protein in the absence of genetic manipulation. dBRD9-A is a highly optimized heterobifunctional ligand that specifically targets the BRD9 bromodomain and bridges it to the Cereblon E3 ubiquitin ligase complex, thereby targeting BRD9 protein for proteasome-mediated degradation^44,45^ (**Figure 3A**). Treatment of A549 cells with 125nM dBRD9-A led to a rapid depletion of BRD9 protein within 6h (**Figure 3B**), and we could confirm the high specificity of this depletion as levels of the closely related BRD7 protein were unaffected (**Figure 3C**). Importantly, and consistent with our ability to generate BRD9-KO A549 cells, BRD9 degradation by dBRD9-A treatment had negligible effects on cell viability (**Supplementary Figure 3A**). Using dBRD9-A, we confirmed that BRD9 depletion leads to a reduction in IFN-α2-induced MxA protein levels in A549 cells (**Figure 3D**), and to an ineffective mounting of IFN-α2-stimulated antiviral responses against IAV (**Figure 3E**). Such defective mounting of antiviral responses was not limited to the inefficient control of IAV, as dBRD9-A treated A549 cells also failed to induce full IFN-α2-mediated antiviral programs against another RNA virus (Vesicular Stomatitis Virus (VSV)), a DNA virus (Herpes Simplex Virus type 1, HSV1), and a retrovirus (Human Immunodeficiency Virus type 1, HIV1) (**Figures 3F-H**). In addition, dBRD9-A treatment of a panel of human cell-lines (including Hep2, Huh-7, Calu-3 and U87MG), murine 3T3, primary-like human MRC-5, and primary undifferentiated human tracheobronchial epithelial (HTBE) cells, revealed that BRD9 contributes to IFN-stimulated antiviral activity in a broad range of cell-types and hosts (**Figures 3I-L & Supplementary Figure 3B-D**). The independent ability of dBRD9-A to recapitulate the results from BRD9-KO cells supports the hypothesis that BRD9 promotes IFN-stimulated gene expression. Furthermore, these data highlight the cell-type independent function of BRD9 in the IFN-mediated antiviral response against different viruses.

**Figure 3.**
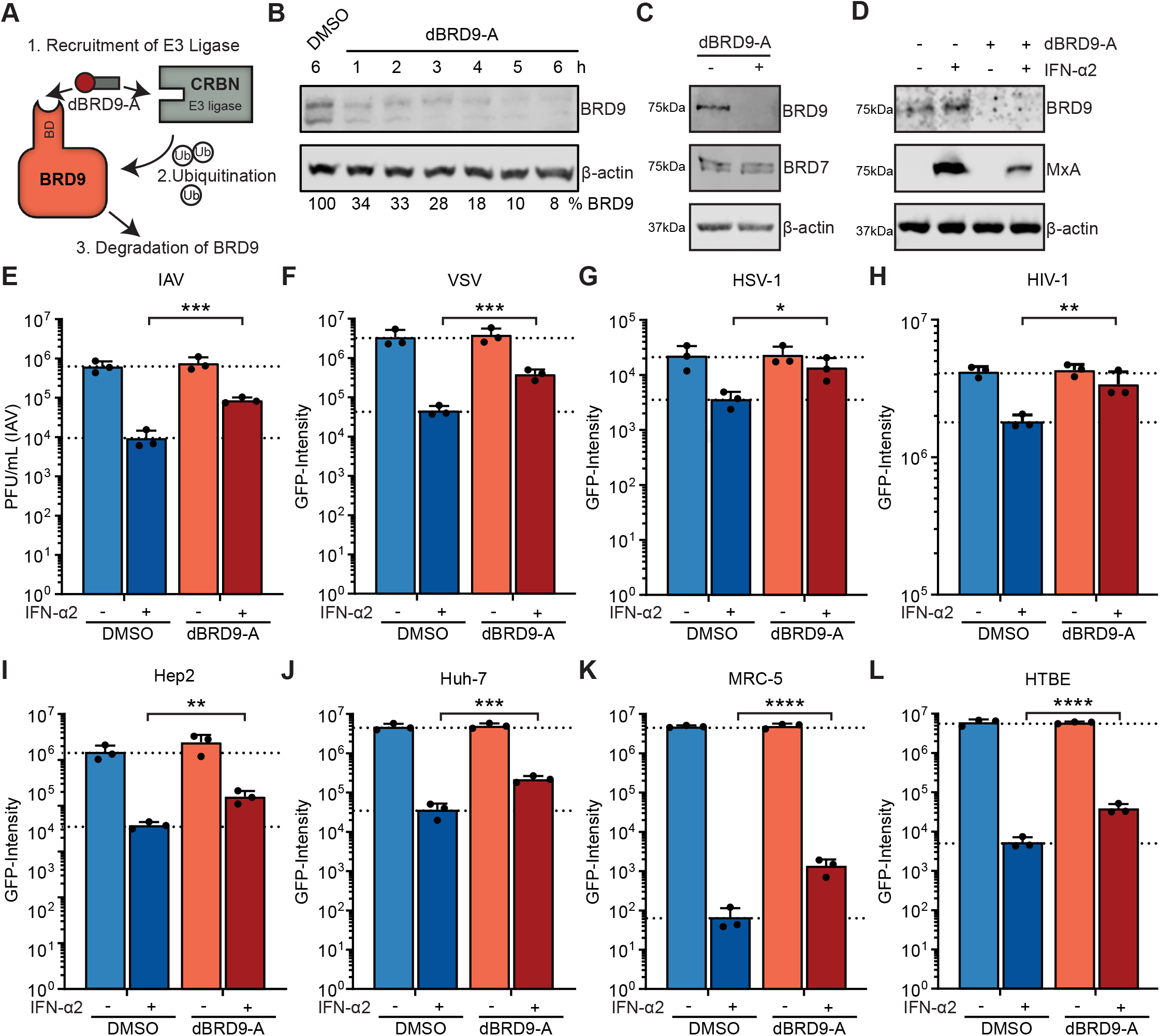
Targeted Degradation of BRD9 Reveals its Cell-Type Independent Contribution to Interferon-Stimulated Antiviral Activity Against Multiple Viruses. **(A)** Schematic representation of the mechanism of action of dBRD9-A. dBRD9-A simultaneously recruits the ubiquitin E3 ligase Cereblon (CRBN) to the bromodomain (BD) of BRD9, thereby facilitating the ubiquitination and degradation of BRD9. **(B)** A549 cells were treated for the indicated time with either DMSO or 125nM dBRD9-A. Following lysis and western blot analysis for the indicated proteins, BRD9 levels were quantified by densitometry and made relative to levels in DMSO-treated cells (% below). Data are representative of at least two independent replicates. **(C)** A549 cells were treated with 125nM dBRD9-A (+) or DMSO (−) for 6h prior to lysis and analysis by western blot for the indicated proteins. Data are representative of at least two independent replicates. **(D)** A549 cells were treated with 125nM dBRD9-A (+) or DMSO (−) for 6h prior stimulation with 1000 IU/mL of IFN-α2 for 16h. Following lysis, western blot was used to detect levels of the indicated proteins. Data are representative of at least two independent replicates. **(E)** A549 cells were treated as described in (D), but following stimulation with 1000 IU/mL of IFN-α2 for 16h were infected with IAV (WSN/33) at an MOI of 0.01 PFU/cell. Viral titers were determined after 24h by plaque assay. Data represent means and standard deviations from three independent experiments (individual data points shown). Statistical significance was determined by 1-way ANOVA on log-transformed plaque counts (***p-value < 0.001). **(F-H)** A549 cells were treated as described in (E), but IFN-α2 concentrations ranged from 10-1000 IU/mL depending upon the virus, and infections were similarly performed but with: VSV-GFP (F); HSV1-GFP (G); or VSV-G pseudotyped HIV1-GFP (H). **(I-L)** Hep2 (I), Huh-7 (J), MRC-5 (K), and undifferentiated HTBE (L) were treated and infected with VSV-GFP as in (F). For F-L, Total Integrated Green Fluorescent Intensities were determined using the Incucyte live-cell analysis system after 10h (G), 24h (F, I-L) or 48h (H). Data represent means and standard deviations from three independent experiments (individual data points shown). Statistical significance was determined by 1-way ANOVA on log-transformed intensity values (*p-value < 0.05; **p-value < 0.01; ***p-value < 0.001; ****p-value < 0.0001). See also **Supplementary Figure 3**.

### BRD9 Promotes the Interferon-Stimulated Expression of a Subset of Antiviral Genes

To gain a deeper, and system-wide understanding of the spectrum of IFN-stimulated genes regulated by BRD9, we conducted a transcriptomic experiment to identify genes differentially-expressed upon IFN-α2 stimulation of A549 cells in the presence or absence of dBRD9-A (**Figure 4A**). Experiments were performed three independent times, and we defined up- or down- regulated genes using cut-off values of 2-fold-change (FC) and a p-value of ≤ 0.001. Using these criteria, we identified 333 significantly upregulated, and 5 significantly downregulated, gene transcripts in A549 cells following 6h of 100 IU/mL IFN-α2 treatment (**Figure 4B & Supplementary Dataset 2**). Notably, dBRD9-A treatment alone resulted in the downregulation of only 11 gene transcripts, which is similar to independent transcriptomic analyses of dBRD9-A action (**Supplementary Dataset 2**). None of the dBRD9-A affected genes have previously been implicated in the type I IFN signaling pathway or have known antiviral activity. Of the 333 IFN-α2-stimulated genes (ISGs) identified in A549 cells, we found that prior dBRD9-A treatment led to significantly reduced induction of 29 ISGs, including many ISGs known to harbor antiviral activity against the viruses used in this study, such as *MX1*, *MX2*, *IFITM1*, *IFITM3*, *IDO1* and *BST2* (**Figure 4C & Supplementary Dataset 2**). Importantly, the expression of many ISGs was not affected by dBRD9-A treatment, and it was notable that the impact of dBRD9-A on ISG expression did not appear to directly correlate (positively or negatively) with gene induction levels by IFN-α2, indicating gene specificity (**Figures 4D-E**). Targeted RT-qPCR analysis confirmed the inhibitory effect of dBRD9-A treatment on IFN-α2-stimulated expression of *MX1*, *MX2* and *IFITM1*, but not *IFIT1* or *ISG15* (**Figure 4F**). Strikingly, this gene-specific pattern was not identical between different human cell-lines (**Supplementary Figure 4A-C**), suggesting that epigenetic factors regulating chromatin state (rather than specific ISG promoter sequences) likely determine the contribution of BRD9 to ISG expression. Indeed, we could not find an enrichment of any specific promoter sequences in the ISGs regulated by dBRD9-A treatment (*data not shown*). These data indicate that BRD9 has a role in promoting the IFN-stimulated expression of a broad set of specific ISGs, and that this set of ISGs may be cell-type, or cell-state, specific.

**Figure 4.**
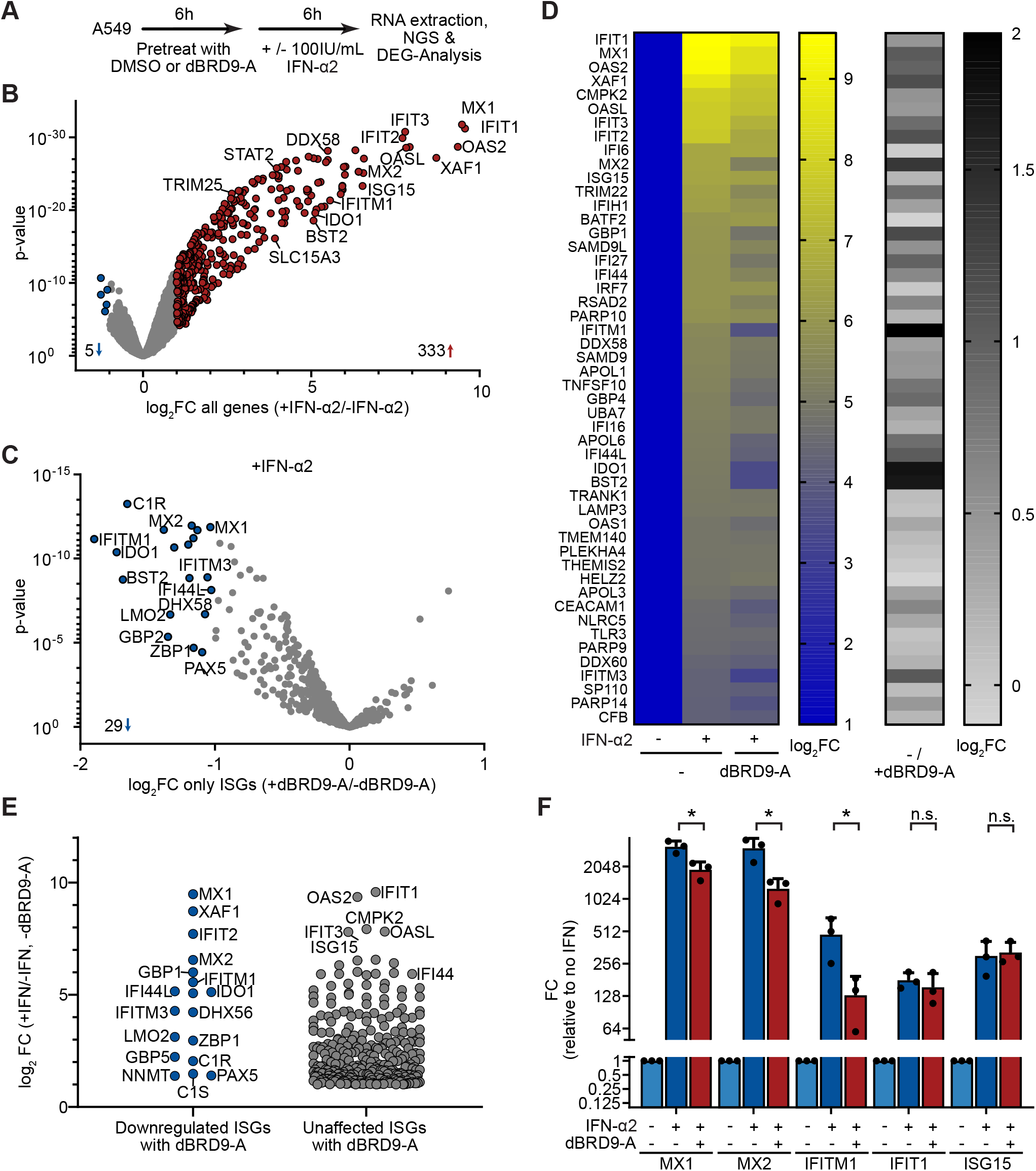
BRD9 Promotes the Interferon-Stimulated Expression of a Subset of Antiviral Genes. **(A)** Overview of the dBRD9-A transcriptomics experiment in A549 cells. **(B)** Up-(red) and down-(blue) regulated genes identified in A549 cells following treatment with 100 IU/mL IFN-α2 for 6h: ISGs were defined as induced by IFN-α2 with FC>2, p-value ≤ 0.001. **(C)** Impact of dBRD9-A pre-treatment on the induction of ISGs identified in (B). Colors indicate IFN-induced transcripts significantly affected (blue) or unaffected (gray) by dBRD9-A pretreatment (FC>2, p-value ≤ 0.001). **(D)** Heatmaps highlighting the gene expression changes observed with each treatment. Left: the top 50 ISGs, ranked by IFN-induced expression (B), were compared with respect to the impact that dBRD9-A pre-treatment had on induction. Right: comparative fold change in induction for each ISG in the presence or absence of dBRD9-A. **(E)** ISG expression level following IFN-α2 stimulation, grouped by whether they were affected or not by dBRD9-A pretreatment (see C). **(F)** RT-qPCR confirmation of transcriptomics results for selected transcripts (*MX1*, *MX2*, *IFITM1*, *ISG15* and *IFIT1*) in A549 cells treated as outlined in (A). *GAPDH* transcript levels were used for normalization. Data represent means and standard deviations from three independent experiments (individual data points shown). Statistical significance was determined by unpaired 2-tailed t-test on ΔC_t_ values (*p-value < 0.05; n.s. not significant). See also **Supplementary Figure 4** and **Supplementary Dataset 2**.

### BRD9 Function Requires its Acetyl-Binding Activity and Unique DUF3512 Scaffolding Domain

BRD9 comprises an N-terminal bromodomain and a C-terminal DUF3512 scaffolding domain that mediates a specific interaction with the GLTSCR1/1L component of ncBAF^24,46^ (**Figure 5A**). Bromodomains are reader-modules of acylated residues, and generally mediate the interaction of protein complexes, such as ncBAF, with DNA via acetylated histones. The bromodomain of BRD9 is reported to exhibit high binding affinity towards acetylated lysines as well as an unconventional binding of butyrylated lysines^47^. To test the impact of BRD9 activities on type I IFN-induced function, we initially constructed a series of BRD9 mutants that either lacked the entire bromodomain (dBD), or which contained single amino-acid substitutions known to abolish the ability of BRD9 to bind acetylated lysines (N216A) or to bind butyrylated lysines (M208I)^47^. Lentivirus-mediated reconstitution of BRD9-KO cells with either wildtype (wt) BRD9 or the individual BRD9 mutants revealed that all constructs were expressed similarly (**Figure 5A**). However, there were clear differences in the ability of each BRD9 construct to restore the antiviral activity of IFN-α2: as compared to wt BRD9, BRD9 mutants lacking the entire bromodomain (dBD) or lacking acetyl-lysine binding (N216A) were inefficient at reconstituting IFN-α2 activity, while the BRD9 mutant lacking butyryl-lysine binding (M208I) functioned similarly to wt BRD9 (**Figure 5B**). These data suggest that acetyl-(but not butyryl-) binding by the BRD9 bromodomain is important, although not essential, for the function of BRD9 in the context of a type I IFN-mediated antiviral response.

**Figure 5.**
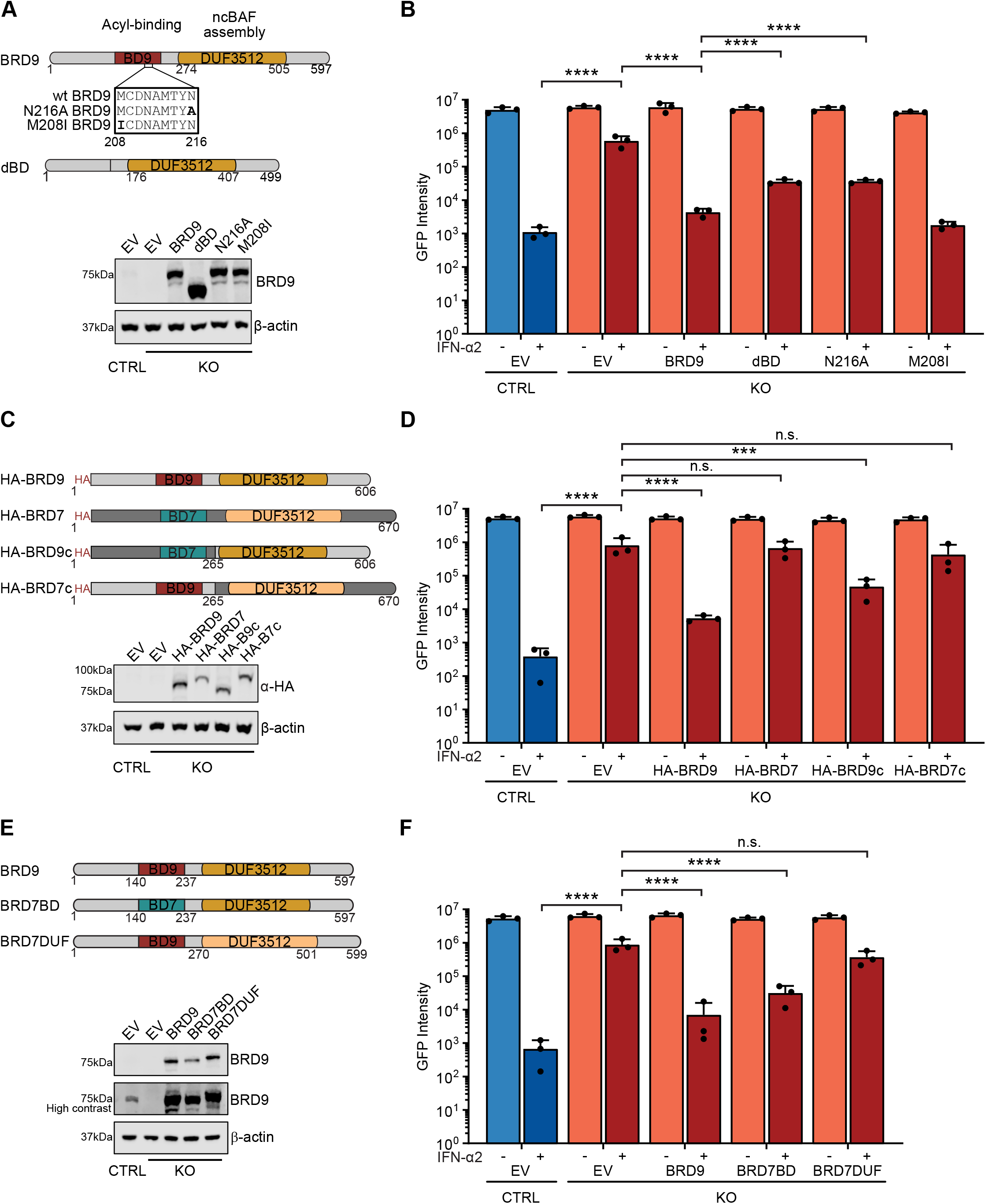
BRD9 Function Requires its Acetyl-Binding Activity and Unique DUF3512 Scaffolding Domain. **(A)** Top: schematic representation of wild-type (wt) human BRD9 and selected mutants, highlighting the acyl-binding bromodomain (BD9) and the DUF3512 domain responsible for ncBAF assembly. Numbers represent amino-acid residues. The box highlights BD9 positions 208-216, which include residues critical for acetyl (N216) or butyryl (M208) binding. dBD is an engineered BRD9 variant lacking the entire bromodomain (residues 140-237). Bottom: western blot analysis of CTRL or BRD9-KO cells stably transduced with lentiviruses expressing empty vector (EV), or wt and mutant BRD9 constructs. **(B)** The various CTRL and BRD9-KO cells described in (A) were stimulated with 1000 IU/mL of IFN-α2 for 16h prior to infection with VSV-GFP at an MOI of 0.6 PFU/cell. Total Integrated Green Fluorescent Intensities were determined using the Incucyte live-cell analysis system at 48h post-infection. **(C)** Top: schematic representation of HA-tagged wt human BRD9, BRD7, and selected chimeric mutants. BD7 is the BRD7 bromodomain. Numbers represent amino-acid residues of the final constructs. Bottom: western blot analysis of CTRL or BRD9-KO cells stably transduced with lentiviruses expressing empty vector (EV), or wt and chimeric mutant constructs. **(D)** The various CTRL and BRD9-KO cells described in (C) were stimulated and infected as described in (B). **(E)** Top: schematic representation of wt human BRD9, and selected chimeric mutants with domains from BRD7. Numbers represent amino-acid residues of the final constructs. Bottom: western blot analysis of CTRL or BRD9-KO cells stably transduced with lentiviruses expressing empty vector (EV), or wt and chimeric constructs. **(F)** The various CTRL and BRD9-KO cells described in (E) were stimulated and infected as described in (B). For (B), (D) and (F), data represent means and standard deviations from three independent experiments (individual data points shown). Statistical significance was determined by 1-way ANOVA on log-transformed intensity values (***p-value < 0.001; ****p-value < 0.0001; n.s. not significant).

We next investigated the contribution of the BRD9 DUF3512 domain to type I IFN-mediated antiviral activity. Highly homologous DUF3512 domains are shared between BRD9 and the closely related BRD7, although each serves as a specific scaffolding interaction platform for either the ncBAF component, GLTSCR1/1L, or the PBAF component, ARID2, respectively^24,46^ (**Supplementary Figure 1B**). We generated N-terminally HA-tagged chimeras between BRD9 and BRD7 (**Figures 5C-D**), as well as specific BRD7/BRD9 domain swap constructs (**Figures 5E-F**), and used lentivirus vectors to reconstitute BRD9-KO cells with each variant. All constructs were expressed similarly to their respective wt BRD9 control, and at levels above endogenous BRD9 levels in unedited CTRL cells (**Figures 5C & E**). As compared to wt BRD9, BRD7 expression was unable to restore the antiviral activity of IFN-α2 in BRD9-KO cells (**Figure 5D**), highlighting the specificity of BRD9 function in this pathway. Notably, constructs consisting of the BRD9 bromodomain in the context of the BRD7 DUF3512 domain also failed to restore the antiviral activity of IFN-α2 in BRD9-KO cells, while constructs containing the BRD9 DUF3512 domain (even in the context of the BRD7 bromodomain) exhibited a strong ability to restore IFN-α2 function, albeit not to levels equivalent to full-length BRD9 (**Figures 5D & F**). These data suggest that acetyl-binding is important for BRD9 function in the type I IFN-mediated antiviral response independently of bromodomain identity. Furthermore, in this antiviral process, BRD9 must act via the specific ncBAF complex, likely using GLTSCR1/1L, as altering the DUF3512-mediated complex composition attenuated BRD9 function in the type I IFN-mediated antiviral response. Such features of general BRD9 function have been observed previously^48^, and it is noteworthy that *GLTSCR1* was enriched in a previous screen to identify positive regulators of type III IFN-stimulated gene expression^8^. These data support the hypothesis of specific involvement of the ncBAF complex in promoting IFN-stimulated gene expression and antiviral activity.

### Interferon Induces the Close Proximity of BRD9 with STAT2

To further understand the involvement of BRD9 (and the ncBAF complex) in IFN-stimulated gene expression, we sought to identify the BRD9 protein interactome in A549 cells, as well as any IFN-induced changes to this interactome. We therefore N-terminally tagged wt BRD9 with the highly active and promiscuous biotin ligase, TurboID^49^, with the aim of biotinylating proteins proximal to BRD9 in living cells and subsequently capturing them with streptavidin beads prior to identification by mass spectrometry. The benefits of using TurboID for detecting protein-protein interactions include the *in situ* nature of biotin labelling reducing the likelihood of identifying artefactual interactions, and the ability to identify weak or transient interactions in the local environment that might otherwise be lost with other affinity-based methods^49^. For these assays, TurboID-tagged mCherry served as a negative control (**Figure 6A**). Using lentivirus vectors, both TurboID-tagged mCherry and TurboID-tagged BRD9 were independently expressed in A549 cells, and their biotinylated interactomes under resting conditions were determined (**Figure 6B**). By comparison with the mCherry negative control across 3 independent replicates, we identified (with high-confidence) 145 proteins in close proximity to BRD9 under these non-stimulated conditions (**Supplementary Dataset 3**). Notable among the interactors were 7 subunits of the ncBAF complex, including GLTSCR1/1L (also known as BICRA/L) (**Figure 6C**). Furthermore, ARID2, a specific interactor of the BRD7 DUF3512 domain and key component of the PBAF complex, was not identified as a BRD9 interactor. Gene ontology (GO) analysis revealed that these 145 protein interactors are functionally enriched for cellular processes including chromatin remodeling/modification, regulation of transcription, and nucleosome/histone activity (**Figure 6D**). These analyses validate the specificity of our approach in identifying proteins in close proximity to BRD9 and the ncBAF complex in A549 cells.

**Figure 6.**
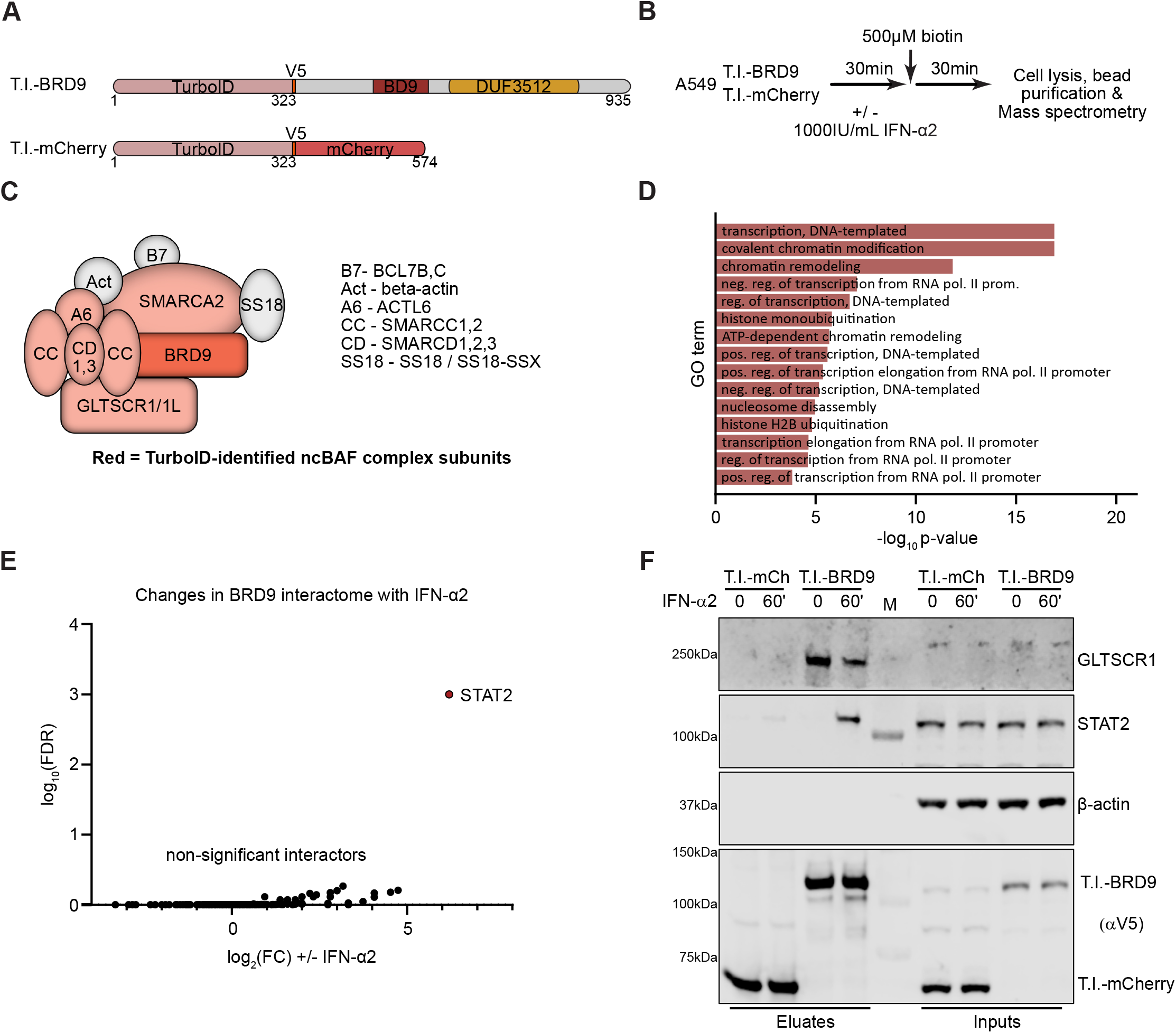
Interferon Induces the Close Proximity of BRD9 with STAT2. **(A)** Schematic representation of TurboID-tagged (T.I.) mCherry (negative control) and TurboID-tagged BRD9. V5 indicates the location of a V5-tag. Numbers represent amino-acid residues of the final constructs. Both constructs were stably expressed in A549 cells. **(B)** Overview of the proximity labelling workflow: interactomes of T.I.-mCherry or T.I.-BRD9 were determined in the presence/absence of a 60min stimulation with 1000 IU/mL of IFN-α2 and 500μM biotin for 30min. **(C)** Schematic of the non-canonical BAF (ncBAF) complex, with factors identified with high confidence (SaintExpress Score ≥ 0.99) by proximity labeling to be specific interactors of BRD9 (as compared to mCherry) colored in red. **(D)** GO term analysis of the 145 interactors identified to be specific to BRD9. **(E)** Comparative BRD9 interactome analysis in the presence/absence of a 60min stimulation with 1000 IU/mL of IFN-α2. Note that FDR for STAT2 was 0, but has been set to 0.0001 for visualization purposes. **(F)** Western blot analysis of samples from independent small-scale streptavidin purifications (termed ‘Eluates’) following the proximity labelling approach outlined in (B). The indicated proteins were detected with specific antibodies. M indicates the marker lane. Data are representative of at least two independent replicates. See also **Supplementary Dataset 3**.

We next used the TurboID assay to study the BRD9 interactome in the presence or absence of a 60-minute stimulation with IFN-α2, which was chosen as a timepoint representing the initiation of ISG transcription. The deduced BRD9 interactomes under these two conditions were essentially identical, except that IFN-α2 stimulation lead to the significant enrichment of STAT2 as a biotinylated factor in the close environment of BRD9 (**Figure 6E & Supplementary Dataset 3**). These results were confirmed in independent proximity labelling, affinity purification and western blot experiments, where GLTSCR1 could be validated as a constitutive and specific interactor of BRD9, while the association of STAT2 with BRD9 was only observed following IFN-α2 treatment (**Figure 6F**). This is reminiscent of a previous observation that the central catalytic BAF subunit, SMARCA4, can directly interact with STAT2, and that this interaction can be enhanced by IFN-α stimulation^37^. Notably, A549 cells appear to express a severely truncated and non-functional SMARCA4^50–52^, and the SMARCA4 paralogue SMARCA2 may compensate for SMARCA4-deficiency in BAF complexes during antiviral responses^52^. Together, these data suggest a functional interaction between BRD9-containing ncBAF complexes and STAT2 following IFN stimulation that likely contributes to driving IFN-stimulated gene expression.

## DISCUSSION

Herein, we describe the identification and characterization of BRD9 as a host factor required for the efficient expression of a subset of ISGs and for the full antiviral activity of IFN. Mechanistically, BRD9 appears to act downstream of cytoplasmic JAK-STAT signaling and STAT protein nuclear translocation, but likely impacts the IFN-induced transcription of ISGs following its proximal association with nuclear STAT2-containing transcription factor complexes (i.e. the ISGF3 complex). BRD9 is a defining member of the ncBAF complex, and its C-terminal DUF3512 domain is essential for recruiting a key component, GLTSCR1/1L, into this complex. We found that switching the recruitment function of BRD9 from GLTSCR1/1L to ARID2 (a PBAF complex component), using a previously-described domain-swap technique^24^, compromised the ability of BRD9 to promote the antiviral effects of IFN, suggesting an important and specific role for the ncBAF complex as a whole in the transcription-promoting activity of IFN. In support of this, we note that *GLTSCR1*, like *BRD9*, was also enriched in a separate screen to identify positive regulators of IFN-stimulated gene expression^8^. While some components of the canonical BAF and PBAF complexes (e.g. SMARCB1/BAF47 and ARID2/BAF200), as well as the central SMARCA4/BRG1 subunit, have previously been implicated in promoting specific ISG transcription in knock-down experiments^22,36–39,53^, no other BAF or PBAF exclusive components were enriched in our genome-wide knock-out screen. This could indicate that these other factors are essential for cell viability, at least in the A549 cell-line used here, or are already non-functional, as is reported for SMARCA4^50–52^. It is also possible that distinct BAF complexes differentially regulate specific ISG subsets in different cell-types depending upon epigenetic factors or subunit expression levels, such that the presence of specific BAF complex compositions within cells actively shapes their individual ISG-profiles in response to IFN. Nevertheless, our findings here indicate that BRD9 is important for the antiviral ISG response (and can be targeted for inhibition) in a range of primary and transformed human (and murine) cells, indicating that at least the ncBAF complex plays an important role in selected ISG expression in different cell-types.

BRD9 function requires an intact bromodomain, and the ability to bind acetylated-lysines is necessary for its promotion of IFN-stimulated antiviral activity. Presumably, this mediates the recruitment of the entire ncBAF complex to a specific target that remains to be elucidated, but which is likely to be acetylated histone tails^47^. Thus, the simplest interpretation of our findings, which is consistent with a previously published model of BAF complex association with the promoters of regulated genes^36^, is that ncBAF constitutively localizes to some key gene promoters (including critical ISGs) via the acetyl-binding property of BRD9, where this complex functions to increase chromatin accessibility and thereby ‘prime’ maximal ISG induction following IFN stimulation (**Figure 7**). We favor this model, as our proximity labelling proteomic assays did not reveal dramatic changes in the BRD9 interactome that might have indicated IFN-stimulated recruitment of BRD9 to new genomic sites. Rather, our data suggest recruitment of the STAT2-containing ISGF3 transcription factor complex to promoters already harboring BRD9 and ncBAF, which is in line with other reports^37^. However, we cannot rule out a more nuanced mechanism. For example, our proteomic experiments confirmed the previously described interaction of BRD4 with the BRD9-containing ncBAF complex^25,54,55^, and BRD4 can mediate the recruitment of ncBAF subunits to chromatin in a bromodomain-dependent manner^55^. Furthermore, BRD4 itself has been implicated in the transcriptional regulation of ISGs^22,39^, and IFN-induced chromatin recruitment mechanisms have been described in this context^37,39^. Thus, the functional interplay between BRD9 and BRD4 remains to be further investigated, perhaps along with additional known targets of the BRD9 bromodomain, such as acetylated RAD54^56^, acetylated CCAR2^57^, and the acetylated vitamin D receptor (VDR)^58^. In this context, we note that VDR also has a proposed role in regulating the JAK-STAT pathway and IFN-mediated STAT1-dependent ISG responses^59^.

**Figure 7.**
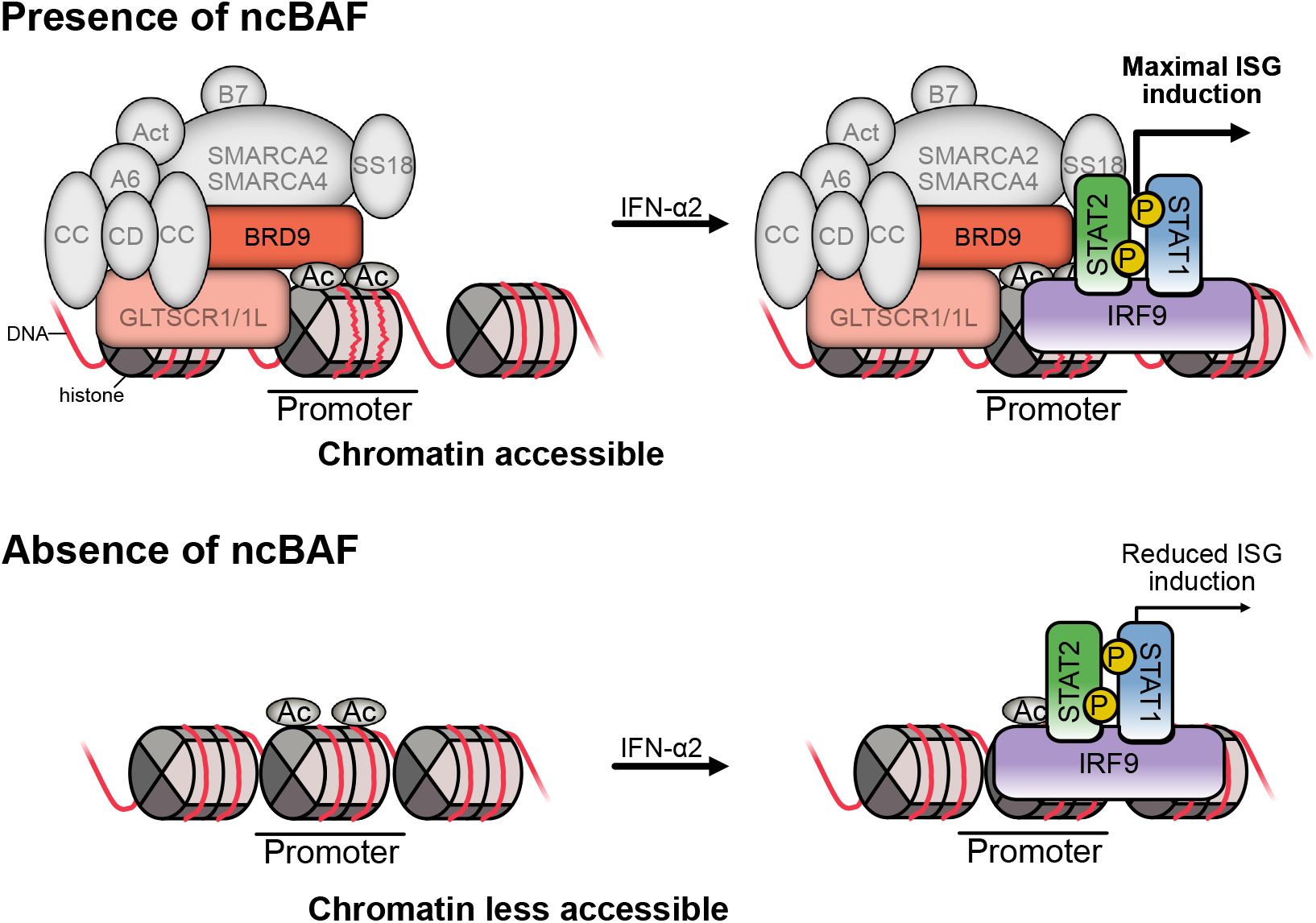
Proposed Model of BRD9 and ncBAF Function at Selected Interferon-Stimulated Gene Promoters. The ncBAF complex is localized to the promoters of regulated genes by the interaction of the BRD9 bromodomain with histone acetylation marks. ncBAF activity maintains the chromatin in an open state, thus enabling the efficient binding of ISGF3 complexes to ISRE sequences following IFN stimulation. Thus, maximal induction of ISG transcription is ‘primed’ to enhance downstream antiviral activity. In the absence of BRD9 (through genetic loss or targeted degradation with small molecules), ncBAF is removed from promoter regions, increasing chromatin compaction and resulting in less efficient transcriptional activation of ISGs following IFN stimulation. This leads to a reduced antiviral activity of IFN.

An important outcome of our study was the observation that small-molecule mediated degradation of BRD9, using dBRD9-A^43,44^, limits IFN-induced expression of certain ISGs in multiple cell-types. This finding could have implications for the consideration of BRD9 as a viable therapeutic target in the treatment of some autoinflammatory interferonopathies^9–17^, particularly under specific circumstances where broad targeting of JAK-mediated signaling may not be appropriate^21,22^. Furthermore, we also note that BRD9 inhibition (or degradation) is an attractive therapeutic aim for the treatment of several cancers^24,44,60^. Therefore, our data should also raise awareness that, in such applications, drugs targeting BRD9 may have the undesirable side-effect of limiting the effectiveness of endogenous host innate antiviral IFN responses, thereby potentially increasing the susceptibility of individuals to some infections.

## METHODS

### Cells and compounds

293T, A549, MDCK, Huh-7, U87MG, Calu-3, Hep2 and mouse 3T3 cells were maintained in Dulbecco’s modified Eagle’s medium (DMEM) (Life Technologies) supplemented with 10% (v/v) fetal calf serum (FCS), 2 mM l-glutamine, 100 units/mL penicillin and 100 μg/mL streptomycin (Gibco Life Technologies). MRC-5 cells were cultured in minimum essential medium Eagle (MEM) supplemented with 10% (v/v) FCS, 100 units/mL penicillin, 100 μg/mL streptomycin, 2 mM l-glutamine (Gibco Life Technologies) and 1% (v/v) non-essential amino acids (Life Technologies). Primary human tracheobronchial epithelial cells were purchased from Lonza and maintained in supplemented Basal medium (Promocell). The original A549/pr(ISRE).eGFP^41^ cells were kindly provided by Rick Randall and Catherine Adamson (University of St Andrews, UK). The A549/pr(ISRE).eGFP.A1 sub-clone was subsequently obtained by limiting dilution of FACS-sorted eGFP-positive cells following 2 rounds of IFN-α2 (Novus Biologicals) stimulation (1000 IU/mL for 16h). The A549/pr(ISRE).eGFP.A1 sub-clone was found to induce high and homogenous levels of eGFP following IFN-α2 stimulation as assessed by the IncuCyte live-cell analysis system (Sartorius) and flow cytometry (see below). Cycloheximide was purchased from Sigma-Aldrich. dBRD9-A^44^ was custom-synthesized by ChemPartner, dissolved in DMSO, and used at the indicated concentration. Cell viability was measured as indicated using the CellTiterGlo (Promega) assay kit, following the manufacturer’s instructions.

### Viruses and infections

Propagation, titration and use of wild-type influenza A virus (IAV; WSN/33 strain) has been described previously^61^. VSV-GFP was a kind gift from Peter Palese (Icahn School of Medicine, New York, USA). HSV1-GFP was kindly provided by Jovan Pavlovic (University of Zurich, Switzerland)^62^. VSV-G pseudotyped HIV1-GFP (NHGΔenvGFP) was generated as previously described using plasmids from Sam Wilson (University of Glasgow, UK)^63,64^. For IAV infections, approximately 1×10^5^ cells were treated or not with 1000 IU/mL of IFN-α2 for 16h prior to infection at the indicated multiplicity of infection (MOI) in PBS supplemented with 0.3% BSA, 1 mM Ca^2+^/Mg^2+^, 100 units/mL penicillin and 100 μg/mL streptomycin. After 1h, infected cells were washed three times with PBS, and then overlaid with DMEM supplemented with 0.1% FBS, 0.3% BSA, 20 mM HEPES, 100 units/mL penicillin, and 100 μg/mL streptomycin. Supernatants were collected at the indicated times post-infection and titrated by standard plaque assay on MDCK cells. For the GFP-encoding reporter viruses (VSV-GFP, HSV1-GFP and pseudotyped HIV1-GFP), approximately 1.5-3×10^4^ cells/well in 96-well plates were treated with DMSO, dBRD9-A and/or IFN-α2 (or universal IFN; PBL Assay Science) as indicated, washed once in PBS, then incubated with the appropriate amount of virus diluted in Fluorobrite DMEM (Life Technologies) medium supplemented with 2% (v/v) FCS, 2 mM L-glutamine, 100 units/mL penicillin and 100 μg/mL streptomycin. Infections were monitored in live cells via GFP expression using the Incucyte live-cell analysis system (Sartorius) for 24-72h. Total Green Integrated Intensity (Green Calibrated Unit x μm²/image) values were exported at the indicated time points.

### CRISPR screening

The full human GeCKOv2 CRISPR knockout pooled library (Addgene #1000000048, a gift from Feng Zhang) was used for genome-wide screening^40^, essentially as described^65^. Briefly, the GeCKOv2 plasmid library with 6 guide RNAs per human gene was propagated in Endura ElectroCompetent cells (Lucigen), and lentivirus stocks were generated by co-transfecting 293T cells with the GeCKOv2 plasmid library, pMD2.G and psPAX2 (Addgene plasmids #12259 and #12260, gifts from Didier Trono) at a ratio of 4:2:1 (GeCKOv2:psPAX2:pMD2.G). After 48h, lentivirus-containing supernatants were clarified by low speed centrifugation and filtration through a 0.45μm filter prior to storage at −80°C. Following titration of the lentiviral library, 1×10^8^ A549/pr(ISRE).eGFP.A1 cells were transduced at a multiplicity of infection (MOI) 0.3 FFU/cell and selected with 1μg/mL puromycin for 14 days. Surviving cells were stimulated with 1000 IU/mL of IFN-α2 (Novus Biologicals) for 16h prior to FACS-based sorting of eGFP-negative cells (Cytometry Facility, University of Zurich), which were then plated into new flasks and allowed to recover. DNA was subsequently isolated from both sorted and non-sorted cells as previously described^66^. Briefly, cells were lysed in 50mM Tris pH 8, 50mM EDTA and 1% SDS, supplemented with proteinase K (Qiagen), and incubated overnight at 55°C. RNA was then digested by addition of RNAseA (Qiagen) for 30min at 37°C. Pre-cooled 7.5M ammonium acetate was added to a final concentration of 1.75M, and remaining impurities were removed by centrifugation. DNA was finally precipitated by addition of isopropanol, washed once in 70% ethanol, and resuspended in a 5mM Tris buffer. Integrated sgRNAs were PCR-amplified by an initial 14 cycles using 7μg/50μL isolated DNA as template. Reactions were pooled, and subjected to a second round of 26 PCR cycles using described primers^65^. All PCRs were performed using Kapa HiFi Hotstart polymerase (Kapa Biosystems). Amplicons were separated on a 2% agarose gel, column-purified (Qiagen) and sequenced on an Illumina NextSeq500 at the Functional Genomics Center Zurich. Data analysis from two independent CRISPR screen experiments, including unsorted control samples, was performed using the PinAPL-Py platform^43^ and the adjusted robust-rank aggregation (αRRA) algorithm with default settings (p-value = 0.01; 1000 permutations).

### Generation of knock-out (KO) cells

Gene knock-outs were generated using either the lentiCRISPRv2 system (Addgene plasmid #52961, a gift from Feng Zhang)^40^ using the methods described above and specific sgRNA-encoding oligos (**Supplementary Dataset 4**), or a ribonucleoprotein (RNP)-based system described previously^67^. Briefly, pre-assembled RNP complexes (see **Supplementary Dataset 4** for crRNA sequences) were delivered into cells by reverse transfection with RNAiMax (ThermoFisher Scientific), and individual cell clones were generated by limiting dilution. Genotypes of selected clones were determined by amplification of the appropriate genomic region and subsequent NGS. Oligo sequences are deposited in **Supplementary Dataset 4**.

### Flow cytometry analysis

A549/pr(ISRE).eGFP.A1 cells were transduced at high MOI (>10 FFU/cell) with a pool of three lentiviruses encoding sgRNAs targeting the indicated factors. After 48h, selection of transduced cells was started with 1 μg/mL puromycin in and maintained for at least 10 days. Transduced cells were then treated with 1000 IU/mL IFN-α2 for 16h, and eGFP expression was quantified in the FITC channel using a BD FACSCanto II.

### Plasmids and lentiviruses

cDNAs encoding BRD7, BRD9, BRD9-dBD and BRD9-N216A were PCR-amplified from existing vectors (Addgene plasmids #65379^68^ and #75114-6^48^; gifts from Kyle Miller and Christopher Vakoc) and cloned into pLVX-IRES-Puro (Takara) via XhoI/NotI or EcoRI/NotI sites. The indicated chimeras of BRD7 and BRD9 were generated by overlapping PCRs or InFusion (Takara). Remaining variants of BRD9 were generated by overlap-extension PCR using Q5 Hot Start High-Fidelity (NEB). TurboID-V5 constructs were generated by first subcloning TurboID from an existing vector (Addgene plasmid #107169^49^, a gift from Alice Ting), together with the V5 linker sequence, into the multiple cloning site of pLVX-IRES-Puro, followed by insertion of BRD9, or mCherry, via EcoRI/NotI sites. All new constructs were authenticated by DNA sequencing. All oligo sequences used for cloning and mutagenesis are available in **Supplementary Dataset 4**. To generate polyclonal cell-lines expressing the gene of interest, lentiviral stocks were initially prepared by co-transfecting 293T cells with each pLVX-IRES-Puro-based plasmid, together with pMD2.G and psPAX2. Lentiviral supernatants were harvested ~60h post-transfection, filtered through a 0.45μm filter, and aliquoted for storage at −80°C. Cells were transduced with the appropriate lentivirus stock for 48h in the presence of 8 μg/mL of polybrene (Millipore) prior to selection with puromycin.

### RT-qPCR analyses

Around 1×10^5^ cells were treated as indicated prior to lysis and RNA isolation using the ReliaPrep™ RNA Miniprep kit (Promega). Approximately 1μg of total RNA was reverse transcribed into cDNA using SuperScript III (Invitrogen) and an oligo(dT) primer (Thermo Fisher). Transcripts were detected using specific primers (**Supplementary Dataset 4**), the Fast EvaGreen qPCR Master Mix (Biotium), and a ABI7300 Real-Time PCR System. Relative gene expression was determined with the ΔΔC_t_ method, using GAPDH for normalization.

### Transcriptome analysis

RNA was extracted as described above and libraries from three independent experiments were prepared and analyzed by the Functional Genomics Center Zurich following the Illumina TruSeq stranded mRNA protocol. Briefly, the quality of RNA and final libraries was first determined using the Agilent 4200 TapeStation System. Libraries were then pooled equimolarily, and sequenced on an Illumina NovaSeq6000 sequencer (single-end 100 bp) with a depth of around 20 million reads per sample. Reads were quality-checked with FastQC. Sequencing adapters were removed with Trimmomatic^69^ and aligned to the reference genome and transcriptome of *Homo sapiens* (GENCODE, GRCh38.p10, release 91) with STAR v2.7.3^70^. Distribution of the reads across genomic isoform expression was quantified using the R package GenomicRanges^71^ from Bioconductor Version 3.10. Minimum mapping quality, as well as minimum feature overlaps, was set to 10. Multi-overlaps were allowed. Differentially expressed transcripts were identified using the R package edgeR^72^ from Bioconductor Version 3.10, using a generalised linear model (glm) regression, a quasi-likelihood (QL) differential expression test, and trimmed means of M-values (TMM) normalization.

### Western blotting

Generally, samples were prepared in urea disruption buffer (6 M urea, 2 M β-mercaptoethanol, and 4% SDS), and DNA was sheared either by passing through a 29G needle or by sonification (Branson 250, 12×0.5/1s pulses at 10% amplitude). For detection of BRD9 in A549 cells, lysates were prepared on ice for 20min in RIPA buffer (50mM Tris-HCl pH8.0, 150mM NaCl, 0.1% SDS, 1% sodium deoxycholate, 1% Triton X-100), DNA sheared as described above, and protein concentration measured by BCA assay (Pierce Thermo Fisher). At least 30μg of protein was loaded onto gels to detect endogenous BRD9. Proteins were separated by SDS-PAGE on NuPAGE Novex or Bolt 4%–12% Bis-Tris gels (Invitrogen) prior to transfer to Protran nitrocellulose membranes (Amersham) at 30V for 90min. Proteins were detected using the following primary antibodies: β-actin (Santa Cruz, sc-47778); MxA (kind gift from Jovan Pavlovic, University of Zurich, Switzerland); FLAG M2 (Sigma, F1804); BRD9 (Bethyl Laboratories, A303-781A-M); BRD7 (Cell Signaling Technology, D9K2T); HA (Cell Signaling Technology, 3724); V5 (Bio-Rad, MCA1360); STAT1 (Cell Signaling Technology, 14994S); pSTAT1 (Cell Signaling Technology, 9167S); STAT2 (Santa Cruz, sc-1668); GLTSCR1 (Santa Cruz, sc-515086); JAK1 (BD Bioscience, 610231); pJAK1 (Cell Signaling Technology, 74129). Signal was detected using the LiCOR system or ECL.

### Immunofluorescence microscopy

5×10^5^ cells were seeded onto cover slips in 12-well plates and treated the next day for the indicated time with 1000 IU/mL IFN-α2, prior to fixation with 3.7% paraformaldehyde (Sigma-Aldrich) for 10 min at room-temperature. Cover slips were washed once in PBS, permeabilized with 1% TritonX-100 (in PBS) for 10 min, then washed a further three times in PBS. Blocking was performed by incubating cover slips for 1h with PBS + 2% BSA (v/v) at room temperature. Following incubation of cells with pSTAT1 and STAT1 antibodies (Cell Signaling 9167S and 14994S), cells were washed three times with PBS before incubation with the appropriate secondary antibody, and a further three washes in PBS. As necessary, DNA was stained by incubating with DAPI (Sigma-Aldrich) for 5min. Following a final three times in PBS and two washes in ddH_2_O, cover slips were mounted onto slides using ProLong Gold Antifade (Life Technologies). Cells were imaged using the Leica DM IL LED Fluo microscope (Leica Microsystems), using the LasX software.

### TurboID proximity labelling

Approximately 5×10^5^ cells stably expressing the indicated TurboID-V5-fusion proteins were treated as described prior to the addition of 500μM biotin (Sigma-Aldrich) for 30min. Cells were subsequently washed five times in PBS, before lysis on ice for 20min in RIPA buffer supplemented with complete protease inhibitors (Roche). DNA was sheared as described above and removed by addition of Benzonase (Millipore) for 1h at 4°C. Lysates were cleared by centrifugation at 16,000 *g* for 10min at 4°C, and a fraction was taken to represent input. The remainder of the sample was processed as described^73^ with slight modifications: briefly, lysates were incubated with streptavidin magnetic beads (Pierce Thermo Fisher) for 1h at room temperature on a rotator to bind biotinylated proteins. Samples were then washed twice in RIPA buffer, once in 1M KCl, once in 0.1M Na_2_CO_3_ and once in freshly prepared 1M urea, 10mM Tris–HCl pH8.0. For proteomic analysis, samples were then washed three times in freshly prepared ABC buffer (50mM ammonium bicarbonate). For western blot analysis, samples were washed three times in RIPA buffer. Proteomic sample preparation and liquid chromatography-mass spectrometry analysis was performed by the Functional Genomics Center Zurich. Proteins were identified using FragPipe (v12.2) and the MSFragger - Ultrafast Proteomics Search Engine^74^. Spectra were searched against a canonical Swiss-Prot *Homo sapiens* proteome database (taxonomy 9606, version from 02/2020), concatenated to its reversed decoyed fasta database. Methionine oxidation was set as variable modification, and enzyme specificity was set to trypsin allowing a maximum of two missed-cleavages. A fragment ion mass tolerance of 0.1 Da and a precursor mass tolerance of 50 PPM were set. The SaintExpress algorithm^75^ was utilized to analyze the spectral count shotgun MS data between samples, with the following settings applied by the CRAPome websuite: lowMode=0; minFold=1; normalize=0.

### Gene ontology (GO) analyses

GO analyses of identified interactors in TurboID experiments, or differential expressed genes in transcriptomics experiment, were performed using medium stringency settings in DAVID^73,74^, using all human genes as background. We excluded BRD9 from the analysis of the proteomics data as it was the bait.

### Statistical analyses

Statistical analyses were performed in GraphPad Prism 9.0.0. FACS MFI data were analyzed by pairwise comparison using an unpaired t-test. Data using virus titers or eGFP intensities were log_10_-transformed and analyzed by unpaired 2-tailed t-test or 1-way ANOVA for comparison of multiple conditions. For RT-qPCR data, ΔC_t_ values were analyzed by unpaired 2-tailed t-test.

## DATA AVAILABILITY

The authors declare that all data supporting the findings of this study are available within the paper and its supplementary information files.

## ACKNOWLEDGEMENTS

We thank Michael Huber, Stefan Schmutz and Marie Pohl (University of Zurich, Switzerland) for helpful discussions and advice relating to practical aspects of this work. We are particularly grateful to Rick Randall and Catherine Adamson (University of St Andrews, UK) for provision of the original A549/pr(ISRE).eGFP cell-line, and to David Remillard (Scripps Research Institute, USA) for advice on dBRD9-A synthesis and use. We also acknowledge Kyle Miller, Peter Palese, Jovan Pavlovic, Alice Ting, Didier Trono, Christopher Vakoc, Sam Wilson and Feng Zhang for providing essential reagents. Amplicon sequencing, transcriptome analyses, and proteome analyses were performed with support from the Functional Genomics Center Zurich. Flow cytometry and FACS were performed with support from the Cytometry Facility, University of Zurich. The research leading to these results received funding from the Swiss National Science Foundation (grant 31003A_182464 to BGH).

## AUTHOR CONTRIBUTIONS

Conceptualization: JB, DE and BGH; Methodology and Investigation: JB, DE, IB, EM, NS, PPP and WWW; Resources: NM and SS; Writing and Visualization: JB, DE and BGH; Supervision: MS and BGH; Funding Acquisition and Project Administration: BGH.

## SUPPLEMENTARY FIGURE LEGENDS

**Supplementary Figure 1. Independent Validation of BRD9 as a Hit in the Genome-Scale Screen for Factors Important for Interferon-Stimulated Gene Expression. (A)** A549/pr(ISRE).eGFP.A1 cells were transduced for at least 10 days with lentiviruses expressing Cas9 and individual sgRNAs derived from the GeCKOv2 CRISPR-Cas9 library targeting BRD9, or STAT1 and IFNLR1. eGFP levels following 16h of IFN-α2 treatment (1000 IU/mL), or mock, were determined by flow cytometry. MFI = mean fluorescence intensity. Data represent means and standard deviations from three independent experiments (individual data points shown). Statistical significance was determined relative to the parental cells stimulated with IFN-α2 using single-tailed ANOVA (*p-value < 0.05; **p-value < 0.01; ***p-value < 0.001; n.s. not significant). **(B)** Schematic representations of the known components of the canonical BRG1- or BRM-associated factors (BAF) complex, the Polybromo-containing BAF (PBAF) complex, and the non-canonical BAF (ncBAF) complex. Factors reported to be unique to each complex are indicated with colored shapes. BRD9 is unique to the ncBAF complex. Related to **Figure 1**.

**Supplementary Figure 2. Data Generated using an Independent Knock-Out Cell Clone to Validate BRD9 as Important for Interferon-Stimulated Antiviral Activity. (A)** An A549-derived BRD9-KO cell-clone (KO#2) was generated using a crRNA targeting exon 1 of *BRD9*. The target sequence of the crRNA (termed crRNA2), and the resulting 1 nt homozygous genomic insertion determined by NGS, are shown in comparison to an unedited control clone (CTRL#2). The generated insertion leads to a premature termination (PMT) codon in the following exon. Encoded amino-acids are shown below the CTRL nucleotide sequence (bold indicates nucleotides in exons). **(B)** Western blot analysis of lysates from CTRL#2 or BRD9-KO#2 cells. BRD9 and β-actin were detected with specific antibodies. Data are representative of at least two independent replicates. **(C)** Western blot analysis of CTRL#2 or BRD9-KO#2 lysates from cells treated for 16h with 1000 IU/mL of IFN-α2. The indicated proteins were detected with specific antibodies. Data are representative of at least two independent replicates. **(D)** CTRL#2 or BRD9-KO#2 cells were treated, or not, with 1000 IU/mL of IFN-α2 for 16h prior to infection with IAV (WSN/33) at an MOI of 0.01 PFU/cell. Viral titers were determined after 24h by plaque assay. **(E)** CTRL#2 or BRD9-KO#2 cells were stably-transduced with BRD9-expressing, or control (EV, empty vector), lentiviruses and treated with a range of IFN-α2 concentrations (0, 10, 100, 1000 IU/mL) for 16h prior to lysis and analysis for the indicated proteins by western blot. Data are representative of at least two independent replicates. **(F)** CTRL#2 or BRD9-KO#2 cells were stably-transduced with BRD9-expressing, or control (EV, empty vector), lentiviruses and treated with 1000 IU/mL of IFN-α2 for 16h prior to infection with IAV (WSN/33) at an MOI of 0.01 PFU/cell. Viral titers were determined after 24h by plaque assay. For (D) and (F), data represent means and standard deviations from three independent experiments (individual data points shown). Statistical significance was determined by 1-way ANOVA on log-transformed plaque counts (*p-value < 0.05; **p-value < 0.01; ***p-value < 0.001). Related to **Figure 2**.

**Supplementary Figure 3. Targeted Degradation of BRD9 is Non-Cytotoxic and Reveals its Cell-Type Independent Contribution to Interferon-Stimulated Antiviral Activity. (A)** A549 cells were treated for 22h with either DMSO, 125nM dBRD9-A, or 1μg/mL cycloheximide (CHX). CellTiterGlo was used to determine cell viability relative to untreated cells. **(B-D)** Calu-3 (B), U87MG (C) or 3T3 (D) cells were treated with 125nM dBRD9-A or DMSO for 6h prior stimulation with 1000 IU/mL of IFN-α2 (B-C) or 400 IU/mL of universal type I IFN (D) for 16h. Cells were then infected with VSV-GFP at an MOI of 0.6 PFU/cell and Total Integrated Green Fluorescent Intensities were determined using the Incucyte live-cell analysis system at 24h post-infection (B-C). Data represent means and standard deviations from three independent experiments (individual data points shown). Statistical significance was determined by 1-way ANOVA on log-transformed intensity values (*p-value < 0.05). For (D), cells were infected with IAV (WSN/33) at an MOI of 0.001 PFU/cell. Viral titers were determined after 52h by plaque assay. Data represent means and standard deviations from three independent experiments (individual data points shown). Statistical significance was determined by 1-way ANOVA on log-transformed plaque counts (**p-value < 0.01). Related to **Figure 3**.

**Supplementary Figure 4. Cell-Type Specific Interferon-Stimulated Gene Subsets are Affected by dBRD9-A Pretreatment. (A-C)** RT-qPCR results of selected transcripts (*MX1*, *MX2*, *IFITM1*, *ISG15* and *IFIT1*) in U87MG cells (A), primary-like MRC-5 cells (B) and undifferentiated HTBE cells (C). Cells were pretreated with 125nM dBRD9-A or DMSO for 6h prior to induction of ISG transcription by addition of 100 IU/mL IFN-α2 for 6h. *GAPDH* transcript levels were used for normalization. Data represent means and standard deviations from three independent experiments (individual data points shown). Statistical significance was determined by unpaired 2-tailed t-test on ΔC_t_ values (*p-value < 0.05, **p-value < 0.01, ***p-value < 0.001, ****p-value < 0.0001, n.s. not significant). Related to **Figure 4**.

## SUPPLEMENTARY DATASETS

**Supplementary Dataset 1.**

Results from the Genome-Scale Loss-of-Function Screen to Identify Human Genes Important for Interferon-Stimulated Gene Expression. Related to **Figure 1**.

**Supplementary Dataset 2.**

Results from the Transcriptomic Analyses to Identify Gene Expression Profiles Altered by dBRD9-A. Related to **Figure 4**.

**Supplementary Dataset 3.**

Results from the Proximity Labeling Proteomic Analyses to Identify Proteins in Close Proximity to BRD9. Related to **Figure 6**.

**Supplementary Dataset 4.**

Table of Oligonucleotides Used in this Study. **Related to Methods**.

