## Supplemental Figure 1 for "Genome-Wide CRISPR Screening Identifies BRD9 as a Druggable Component of Interferon-Stimulated Gene Expression and Antiviral Activity"

### Supplementary Figure 1

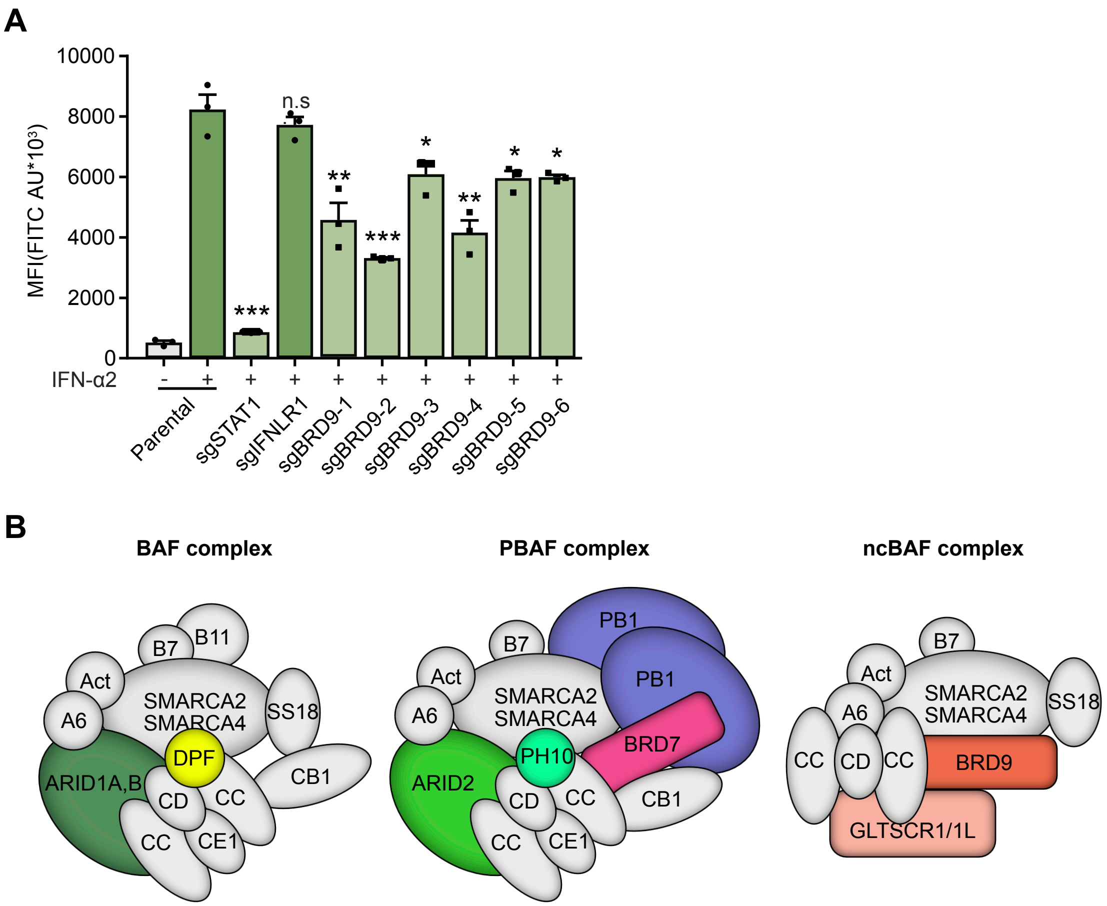


B7 - BCL7B,C
B11 - BCL11A,B
Act - β-actin

CC - SMARCC1/2
CE1 - SMARCE1
DPF - DPF1,2,3

A6 - ACTL6
SS18 - SS18/SS18-SSX
PH10 - PHF10

CD - SMARCD1,2,3
CB1 - SMARCB1
