## Supplementary figures and images for "Genome-Wide CRISPR Screening Identifies BRD9 as a Druggable Component of Interferon-Stimulated Gene Expression and Antiviral Activity"

### Supplemental Figure 2

#### Supplementary Figure 2

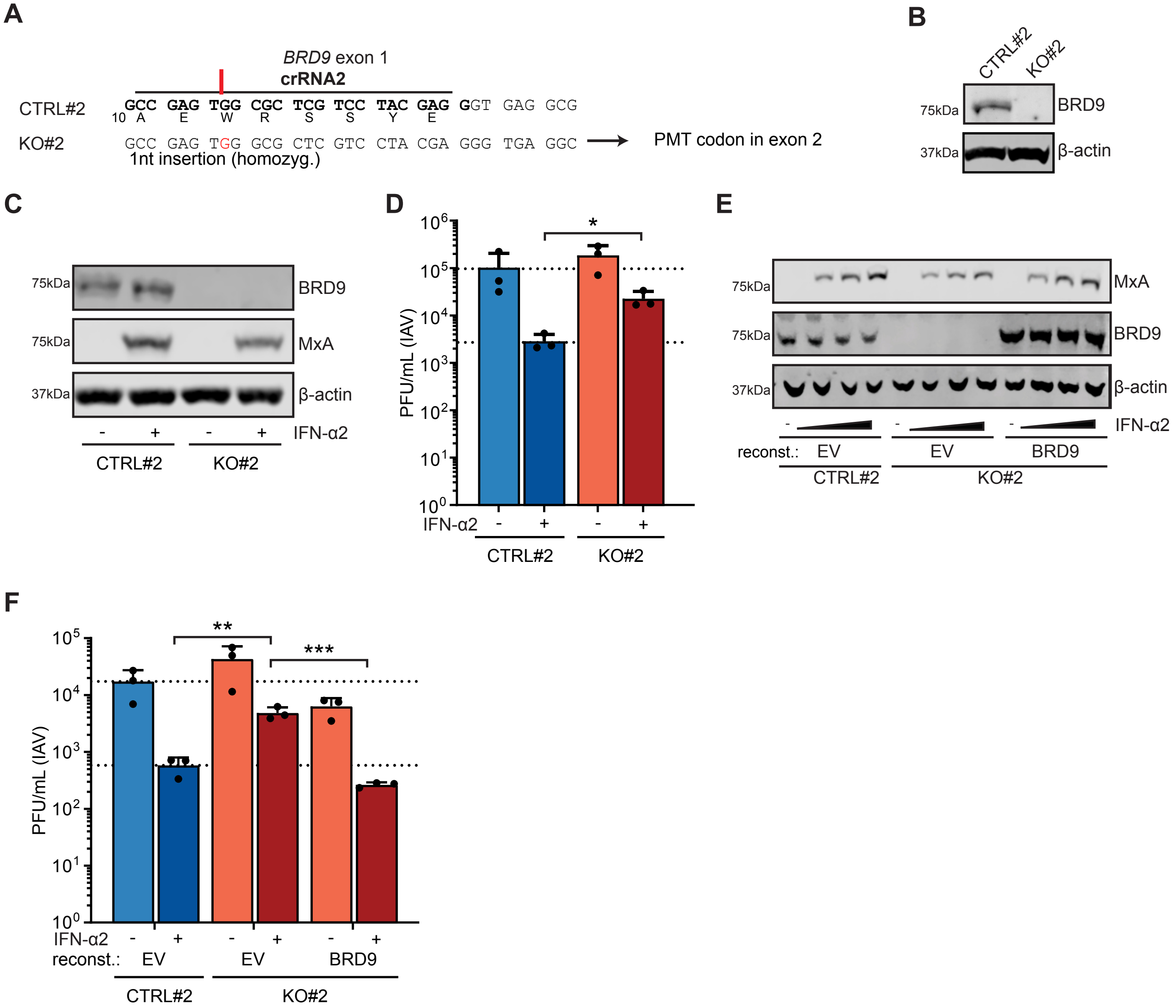

### Supplemental Figure 3

#### Supplementary Figure 3

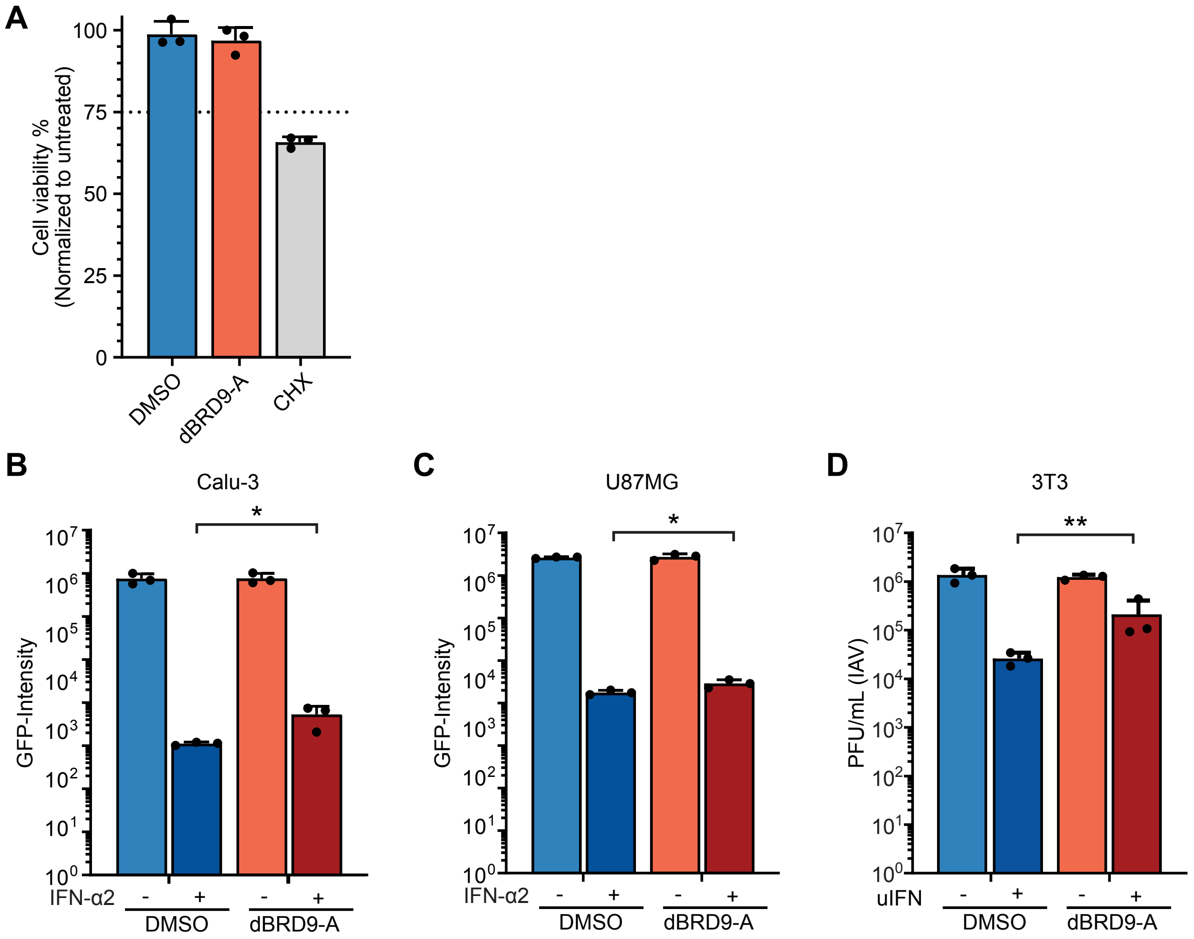

### Supplemental Figure 4

#### Supplementary Figure 4

A

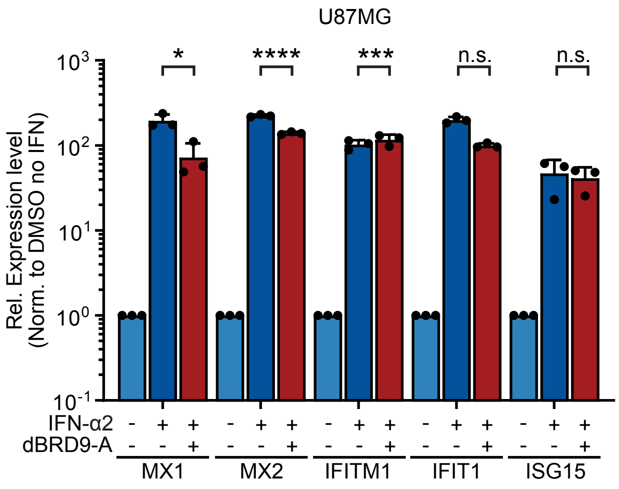


B

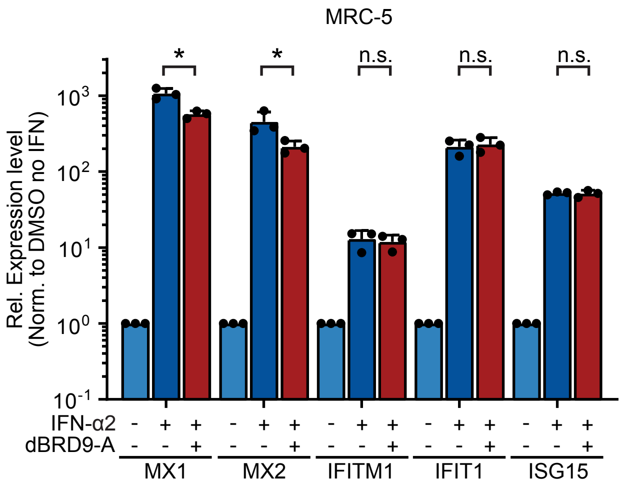


C

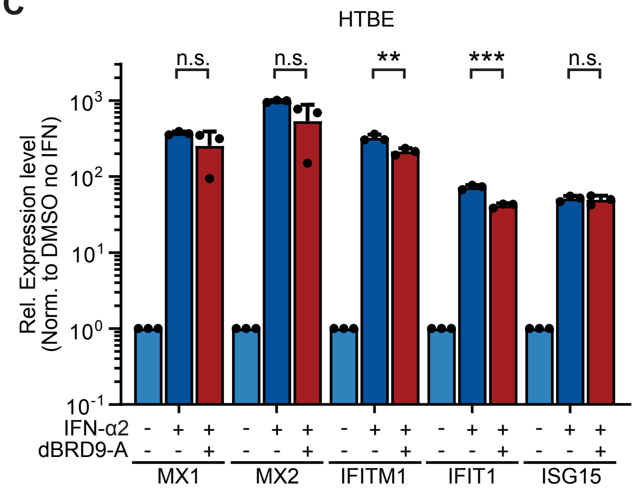
